# The Effects of Phosphorylation on the Structure and Function of Motif A, an Intrinsically Disordered Region within SIRT1

**DOI:** 10.64898/2026.04.16.718858

**Authors:** Sabrina M. Richter, Hoang-Long Bui, Addison Chen, Chloe Tannous, Benjamin R. Butler, Sophia D. Bennett, Samson Quang-anh Nguyen, Justin Prado, Ayan Mohamed, Isaac A. DuBois, Emile Tadros, Nhu Tam Thai, Selina Lima Guan, Carla Marie Peralta, Andy Kwong, Laura M. L. Hawk, Gianmarc Grazioli, Ningkun Wang

## Abstract

The NAD^+^ dependent deacetylase sirtuin-1 (SIRT1) is known to elicit cellular defenses against aging, cancer, and other aberrant pathologies. Previous studies have identified an intrinsically disordered region of SIRT1 comprised of N-terminal residues 1-52, herein referred to as motif A, which activates SIRT1 activity, likely through intramolecular interactions. Additionally, phosphorylation of N-terminal residues Ser27 and Ser47 has been shown to be important for regulating SIRT1 activity and stability. The lack of *in vitro* characterization of these effects hampers our further understanding of the role of motif A in SIRT1 regulation. In this study, we elucidate the role phosphorylation plays in motif A’s structure as well as its regulatory effects on SIRT1 activity against Ac-p65. We find that phosphomimetic mutation at Ser27 significantly increases the activation effect of motif A towards SIRT1. This result is supported by molecular dynamics simulations of the phosphomimetics, which reveal stabilization of different transient structures for motif A depending on whether Ser27 and Ser47 have been modified. A key finding suggested by this study is that phosphorylation of S27 appears to activate SIRT1 by causing motif A, which is intrinsically disordered in the WT, to fold into an ordered structure. This conclusion is based on both the experimental findings and simulation results. These findings contribute to our understanding of SIRT1 regulation, specifically the role played by phosphorylation within the N-terminal disordered region.

## Introduction

The lysine deacetylase SIRT1 is a transcriptional regulator affecting the acetylation state of chromatin proteins, transcription factors, and other regulatory proteins. Well-characterized substrates of SIRT1 include the tumor suppressor protein p53, the metabolic regulator PCG-1α, and the p65 subunit of NF-κB, an activator of inflammatory signaling pathways^1–4^. Deacetylation of these and other targets has implicated SIRT1 involvement in lipid and glucose metabolism, neurodegenerative protection, and cell survival^5^. The promiscuity of SIRT1 underscores its physiological significance and makes it an attractive therapeutic target. It also implies that a complex regulatory scaffolding exists to mediate the activity level of the enzyme in response to distinct environmental signals.

SIRT1 shares a conserved catalytic core with other sirtuin homologs, including the human sirtuins SIRT2-7^6–8^ as well as Sir2, the yeast homolog of SIRT1^9^. Unlike other sirtuins, SIRT1 is distinguished by its extensive N- and C-terminal regions, which interact with the enzyme intramolecularly and potentiate regulatory activities^10,11^. The STAC (sirtuin activating compound) binding domain (SBD), comprised of residues 185-230^8,12^ has been identified as the binding site for small molecule STACs as well as the endogenous nuclear proteins such as active regulator of SIRT1 (AROS) and deleted in breast cancer-1 (DBC-1)^13,14^. However, existing research provides conflicting information on the mechanism of these regulators and their cellular outcomes. Our understanding of the mechanisms of SIRT1 regulation is thereby incomplete, and additional regulatory features may likely exist in the N-terminus which are yet to be characterized.

This study is focused on an intrinsically disordered region with helical tendencies within the N-terminus of SIRT1, coined motif A. Motif A (SIRT1 1-52) has been shown to enhance the enzyme’s deacetylase activity towards acetylated p65, a member of the NF-κB family, and promote SIRT1-mediated glucose tolerance in mice^15^. This regulatory effect is likely due to intramolecular protein-protein interactions between motif A and the rest of SIRT1^14,15^, but the mechanistic details underlying these interactions are not well understood. Additionally, the motif A region of SIRT1 is abundantly phosphorylated^16,17^. Phosphorylation at residues Ser27 and Ser47 is the best studied of these modification sites and is the target of multiple kinases, including mTORC1, JNK1, JNK2, Cdk5, and CaMKKβ^18–22^. The effects of phosphorylation at each of these positions are distinct and tissue-specific. Phosphorylation at S27 is highly variable between cell types and changes the deacetylase activity of SIRT1 in a substrate-dependent manner, whereas phosphorylation of S47 has been reported to decrease deacetylase activity of SIRT1 and controls substrate selection of the enzyme by confining it to the nucleus (residues 30-40 are predicted to be a Nuclear Localization Sequence^23^). However, the molecular underpinnings of these phosphorylation events are poorly understood.

This study aims to understand how phosphorylation at S27 and S47 alters the structure and regulatory function of motif A. Enzyme activity assays revealed how motif A regulates SIRT1 activity differently in its different phosphorylation or phosphomimetic states. We also used circular dichroism, limited proteolysis, and molecular dynamics simulations to identify structural and dynamic changes within motif A caused by an S to D phosphomimetic mutation. This information may serve to better outline the structural and physical parameters underlying a significant and complex mechanism of SIRT1 regulation.

## Materials and Methods

### Cloning of Motif A Phosphomimetic Constructs

pET-28a-His6-TEV-SIRT1(1-52) plasmid was used for generating phosphomimetic mutations at S27, S47 and both S27 and S47. Site-directed mutagenesis (SDM) was performed using the New England Biolabs Q5 SDM kit (Ipswitch, MA). The primers used are shown in the table below. PCR products were transformed into NEB-10β cells, and mutations were confirmed by sequencing. Additional details in Supporting Information.

### Expression and Purification of Protein Constructs

The pET-28a-His6-TEV-SIRT1(1-52) plasmids containing N-terminal His6 tagged SIRT1(1-52) constructs and pET28-smt3-hSIRT1-143 plasmid containing SIRT1-143 were expressed by BL21 (DE3) *E. coli* cells. Motif A constructs were purified by Ni-NTA affinity purification, and SIRT1-143 was purified by Ni-NTA affinity purification and Size Exclusion Chromatography. Protein purity was verified by SDS-PAGE to be more than 85% pure. The concentration of proteins was determined by the Bradford assay. Additional details in Supporting Information.

### Peptides Derived from Motif A (SIRT1(1-52))

Five different peptides spanning the first 52 residues of SIRT1, some with the phosphorylated S27 or S47 residues, were obtained. The names and sequences are listed in Supporting Information. Pep(15-41) and Pep(15-41)S27P were synthesized by the Hawk lab at Grand Valley State University. Pep(1-14), Pep(33-52), and Pep(33-52)S47P were purchased from Elim Biopharm (Hayward, CA). All peptides were prepared to 95% purity as confirmed by HPLC. Additional details of peptide synthesis are in supporting information.

### Circular Dichroism of Motif A Proteins and Peptides

Motif A proteins and peptides were thawed and dialyzed into CD Buffer (10 mM phosphate pH 8.0, 50 mM NaCl, 1 mM DTT) overnight. Concentration of the protein in CD Buffer was determined via Bradford Assay. CD spectra were collected on Jasco-1500 Circular Dichroism Spectrometer by continuous scanning over 195 to 280 nm at 25 °C in a 0.1 cm path length quartz cuvette for all samples. The wavelength step was 0.1 nm with an averaging time of 2 seconds and a scanning speed of 100 nm/min. For each sample, 3 continuous scans were collected and averaged. 3 blank scans were collected by scanning the CD buffer. These blanks were averaged and subtracted from the averaged samples. Molar ellipticity (mean residue ellipticity) was calculated as follows:

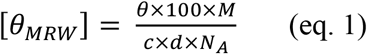

where M is the protein molecular weight, c is the protein concentration in mg/mL, d is the path length in cm, and NA is the number of amino acids per protein.

### Limited Proteolysis of Motif A Constructs

Motif A constructs were divided into two samples: a control sample (no trypsin) and a motif A sample subjected to trypsin digest. Trypsin was added to 10 μM of motif A samples in 50mM HEPES (pH 8.0), 150 mM NaCl, 1 mM CaCl_2_, 1 mM DTT at an E:S molar ratio of 1:2000 and incubated for 0, 15, 30, 45, and 60 min at 37 °C. The same volume of water was added to the control samples. Digestion reactions were stopped at 15-minute intervals by heating samples at 95 °C for 5 min along with adding SDS-PAGE loading dye. Control samples underwent the same procedure. All samples were run on SDS-PAGE and stained with Coomassie Blue.

### Enzyme-Coupled Deacetylation Assay

Deacetylase activity of SIRT1-143 was determined by continuous coupled microplate assay, as previously described^24^. The 13-mer peptide resembling a section of p65 ( RTYETFK(Ac)SIMKKS) was purchased from Elim Biopharm (Harward, CA) and was used as a substrate of SIRT1-143. To determine reaction rates, consumption of NADPH was measured spectrophotometrically at 340 nm. Reactions were performed at 100 μL volumes in a transparent 96 well plate using 640μM NAD^+^, 3.3 mM α-ketoglutarate, 0.2 mM NADPH, 2 μM nicotinamidase PncA, 1 mM DTT, and either 2.5μM or 5μM motif A protein or motif A derived peptide in 20mM phosphate buffer at pH 7.4, with variable substrate concentrations from 2.5-320μM. Reactions were initiated with the addition of either 0.25μM or 0.5μM SIRT1-143 and monitored continuously for 10 minutes at 25°C on a BioTek™ Synergy HTX Multimode Reader. Reaction rates at each substrate concentration were calculated from the linear portion of the reaction using the extinction coefficient for NADPH, 6220 M^-1^cm^-1^, and corrected for basal fluctuation in NADPH levels found in a substrate blank. Reaction rates at each substrate concentration were plotted in GraphPad Prism and fitted to the Michaelis-Menten equation. All measurements were performed in triplicate.

### Molecular Dynamics Simulations

The crystallographic information (CIF) structure files of the different variants of Motif A as well as various segments of Motif A were produced using AlphaFold 3^25^. These initial structure were then inputted into CHARMM GUI, a web-based platform for interactively building preparing input files for molecular dynamics simulations using well-established and reproducible simulation protocols ^26,27^. In this case, CHARMM GUI was used to solvate the proteins using the TIP3P explicit water model in a simulated volume with periodic boundary conditions in order to perform molecular dynamics (MD) simulations using the Nanoscale Molecular Dynamics (NAMD) molecular dynamics software package^28–30^. Each protein was solvated at a pH of 7.5, with ionic strength of 0.15M using K+ and Cl-ions, and the temperature was set to 298K. The equilibration was carried out under NVT ensemble conditions and the production dynamics were run using the NPT ensemble, which is standard practice for stabilizing temperature before engaging the barostat. All MD simulations were run with a standard 2 fs timestep. All molecular dynamics simulations of the proteins were then carried out using NAMD^29,30^, with equilibration runs 200 ps long and production runs carried out for 1.67 *μ*s.

Post-processing of the simulation data involving analysis of interatomic distances was carried out using custom Python codes that leveraged MDAnalysis^31,32^, a Python library for processing molecular simulation data. Additionally, custom TCL scripts were used to harness VMD’s capabilities for identifying secondary structure to produce plots showing secondary structure changing over time across all residues^33^. Molecular visualizations were also carried out using VMD. Additional Python libraries were also used in custom Python scripts for data processing and visualization, including NumPy, Matplotlib, and pandas ^34–36^.

## Results

### Effect of Phosphomimetic Mutations on Motif A Conformation

Motif A encompasses the first 52 amino acids of human SIRT1. This region is rich in proline and glycine, as well as charged residues such as arginine, glutamate, and aspartate. AlphaFold 3^25^ predicts the region to be mostly unstructured with short α-helical stretches (**Figure 1A**). AIUPred^37^ disorder prediction webserver suggests that the motif A region of SIRT1, residues 1-52, is disordered, with the likelihood of a binding region that undergoes a disorder-to-order transition predicted by anchor2^38^ (**Figure 1B**).

**Figure 1.**
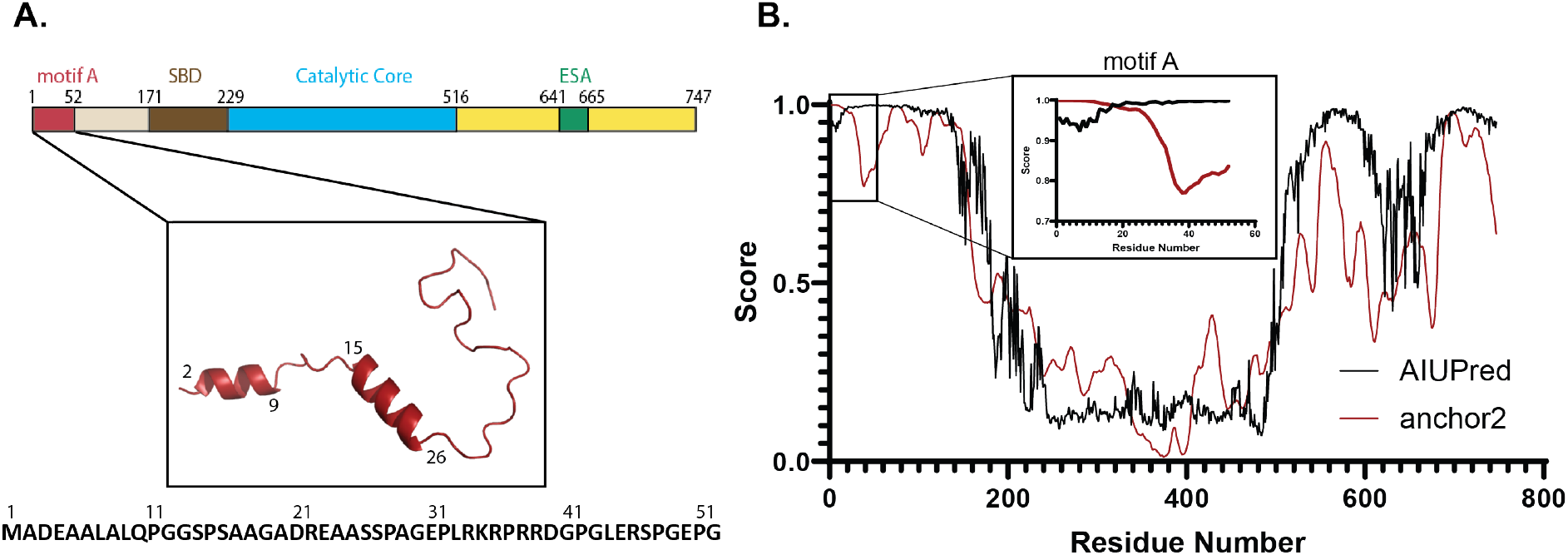
**A)** Schematic showing the position of motif A within SIRT1. SBD stands for the STAC-binding domain, and ESA stands for Essential for Sirtuin Activation. The predicted structure of motif A by AlphFold 3 and the sequence of motif A is also shown. **B)** AIUPred and anchor2 predictions for the sequence of SIRT1. Motif A residues 1-52 are highlighted in the insert box.

To mimic the steric and electrostatic effects of phosphorylation, we introduced the phosphomimetic Ser→Asp mutation at S27, S47 and both S27 and S47 within the motif A protein (SIRT1 residues 1-52). Circular dichroism spectroscopy was obtained for all four different motif A constructs (**Figure 2A**). WT motif A and the motif A S27D, motif A S27D S47D constructs show spectra that are typical for mostly disordered proteins with some α-helical content with characteristic minima for α-helix formation at 208 and 222 nm. Motif A S47D, however, exhibits negative molar ellipticity at the 200 nm region instead of positive ellipticity shown in the other constructs, suggesting increased disorder.

**Figure 2.**
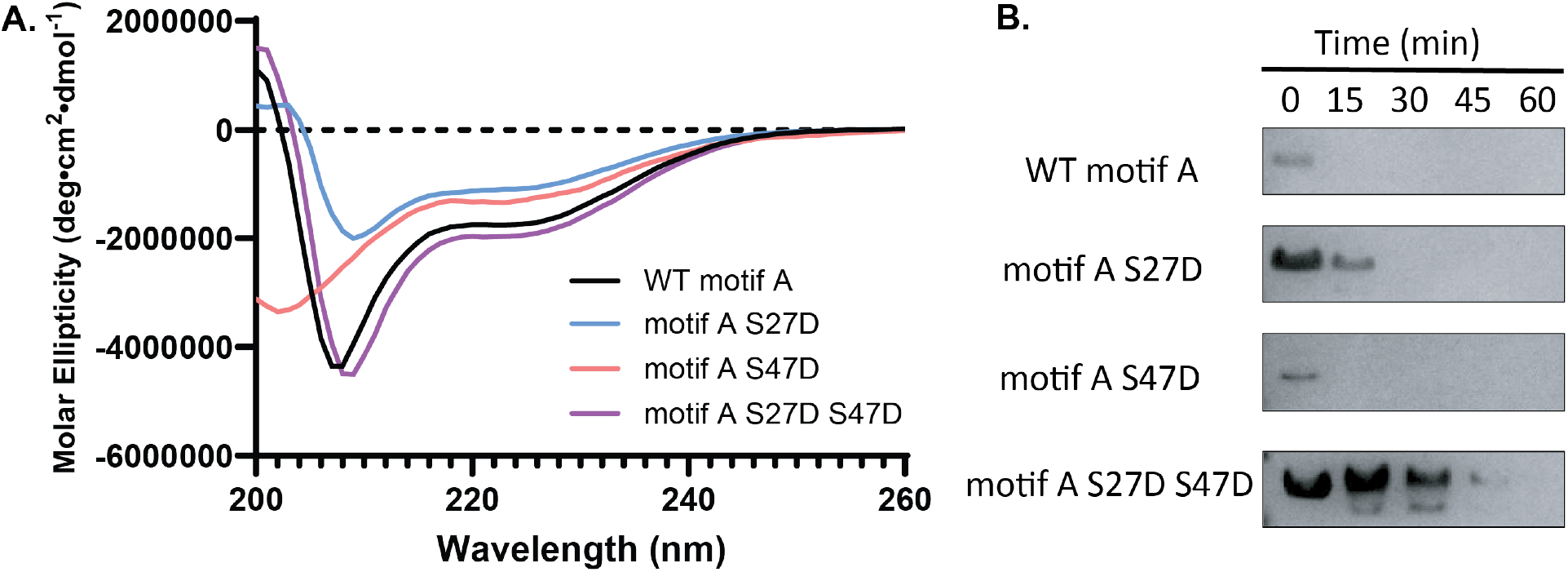
**A)** Overlay of CD spectra for WT motif A, motif A S27D, motif A S47D and motif A S27D S47D. **B)** SDS-PAGE Coomassie gel of limited proteolysis of different motif A constructs with trypsin added at a 1:2000 molar ratio over the course of 60 minutes.

Limited proteolysis experiments on the motif A constructs with 1:2000 trypsin to protein ratio also showed varying degrees of resistance towards trypsin proteolysis. While WT motif A and motif A S47D were immediately degraded by trypsin, motif A S27D and motif A S27D S47D had more resistance to proteolysis, suggesting more folded or “buried” sections within the protein. Notably, motif A S27D S47D exhibited an obvious smaller fragment in the beginning of limited trypsin digestion (**Figure 2B** and **Figure SI 2**).

These results suggest that the phosphomimetic mutations have varying effects on the conformation and stability of motif A.

### Effects of Phosphomimetic Mutations on Motif A Regulation of SIRT1 Deacetylase Activity

To determine how phosphorylation of motif A changes its effect on the enzymatic activity of SIRT1, we added motif A protein (SIRT1 residues 1-52) to a truncated SIRT1-143 construct lacking the N-terminal region and sections of the C-terminal region. SIRT1-143 has previously been characterized by X-ray crystallography and kinetic assays and shows similar activity to full-length SIRT1^7^. We characterized the steady-state kinetics of SIRT1-143 towards an acetylated p65 peptide using a continuous enzyme-coupled assay as previously described^24^, and different constructs of motif A were added to the reaction. Our results (**Figure 3** and **Table 1**) showed that wild-type motif A did not affect the deacetylase activity of the enzyme towards the peptide substrate Ac-p65. Motif A S27D, however, caused a favorable change in substrate recognition of SIRT1-143, lowering the *K*_M_ from 23 ± 3 μM to 14 ± 3 μM, leading to an overall two-fold increase in the specific activity of the enzyme from 900 ± 100 M^-1^s^-1^ to 2000 ± 400 M^-1^s^-1^. Motif A S47D also increased the specific activity of SIRT1-143 to a lesser extent, from 900 ± 100 M^-1^s^-1^ to 1700 ± 200 M^-1^s^-1^, mainly through an increase in *k*_cat_. But this change was not statistically significant (*p*=0.06). The doubly mutated motif A S27D S47D construct altered the kinetic landscape of the enzyme with an increase in *k*_cat_ from 0.021 ± 0.001 s^-1^ to 0.032 ± 0.001 s^-1^; however, the resultant increase in overall efficiency of SIRT1-143 was not statistically significant.

**Figure 3.**
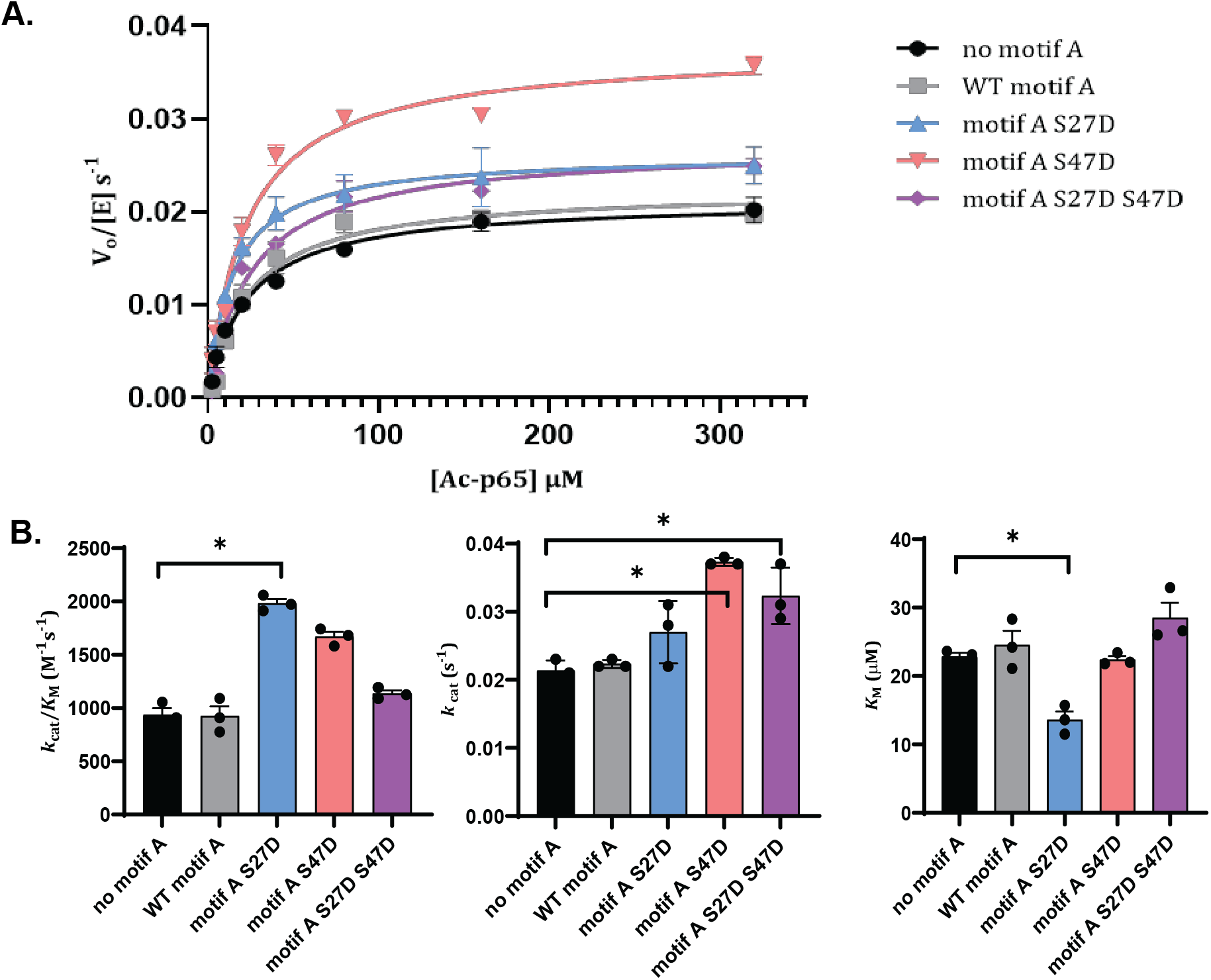
**A)** Enzyme kinetics curves for SIRT1-143 activity against Ac-p65 with and without the addition of various motif A constructs. B) Michaelis-Menten parameters for SIRT1-143 activity with and without the addition of motif A constructs. All kinetics data were collected in at least triplicates and fit with GraphPad Prism. The average and SEM are reported. ^*^ represents *p*-value < 0.05.

**Table 1.** Michaelis-Menten parameters for SIRT1-143 activity with and without the addition of motif A constructs. All kinetics data were collected in at least triplicates and fit with GraphPad Prism. The average and SEM are reported.

| Additive | $k_{\text{cat}} (\text{s}^{-1})$ | $K_{\text{M}} (\mu\text{M})$ | $k_{\text{cat}}/K_{\text{M}} (\text{M}^{-1}\text{s}^{-1})$ |
| --- | --- | --- | --- |
| None | $0.021 \pm 0.001$ | $23 \pm 3$ | $900 \pm 100$ |
| WT motif A | $0.022 \pm 0.001$ | $24 \pm 4$ | $900 \pm 200$ |
| Motif A S27D | $0.027 \pm 0.001$ | $14 \pm 3$ | $2000 \pm 400$ |
| Motif A S47D | $0.038 \pm 0.001$ | $23 \pm 3$ | $1700 \pm 200$ |
| Motif A S27D S47D | $0.027 \pm 0.001$ | $22 \pm 3$ | $1200 \pm 100$ |

### Effects of Phosphorylation on Motif A-derived Peptide Regulation of SIRT1 Deacetylase Activity

To compare whether S to D mutations were sufficient to mimic phosphorylation events within motif A, we chose to also test peptide fragments with and without phosphorylation derived from SIRT1 residues 1-52 that would cover the entire region. While peptide fragments would likely lack global structural characteristics, they would provide valuable information on the local effects near the phosphorylation sites. AlphaFold 3^25^ predicted three helical regions within motif A: residues 3-8, 15-25, and 41-45. Molecular dynamics simulations initiated from these predicted structures showed that not all of these helical regions were persistent throughout the simulations; further, their persistence varied across the 4 different phosphomimetics (more on this in the following section). In addition, MoRFpred^39^ predicted three possible molecular recognition features (MoRFs) within motif A, residues 38-41, 44-45, and 49-50. To best preserve secondary structure and recognition features, we broke down motif A into three overlapping peptide fragments: residues 1-14 (Pep 1-14), containing a predicted α-helix; residues 15-41 (Pep 15-41), containing two predicted α-helices and one predicted MoRF region; and residues 33-52 (Pep 33-52), containing one predicted α-helix and three predicted MoRF regions. Two separate phosphorylated peptides were also prepared: Pep (15-41) S27^P^ and Pep (33-52) S47^P^. CD analysis of all peptides showed typical spectra for disordered peptides (**Figure SI 3**).

We characterized the steady-state kinetics of SIRT1-143 towards an acetylated p65 peptide using a continuous enzyme-coupled assay as previously described^24^, and different motif A-derived peptides were added to the reaction (**Figure 4** and **Table 2**). Unlike what we observed with the motif A protein constructs, all of the peptides positively affected the substrate recognition of the enzyme in nearly identical magnitude, causing a two-fold decrease in *K*_*M*_. This might be due to the fact that the smaller peptide fragments we used eliminated a certain extent of global folding and through-space interactions, which would not be representative of physiological conditions. Regardless, the results suggest that the effects of phosphorylation at each position were distinct, similar to what we observed with motif A proteins. Phosphorylation at position S27 enhanced the activating effect of the peptide on SIRT1-143 deacetylase activity, increasing the specific activity from 1900 ± 300 M^-1^s^-^1 to 3400 ± 500 M^-1^s-^1^, mainly through a decrease in *K*_M_ values. On the contrary, phosphorylation at position S47 decreased the activating effect of the peptide on SIRT1-143 in a small but statistically significant way, from 2200 ± 300 M^-1^s^-1^ to 1700 ± 200 M^-1^s^-1^, mainly through a decrease in *k*_cat_ values.

**Figure 4.**
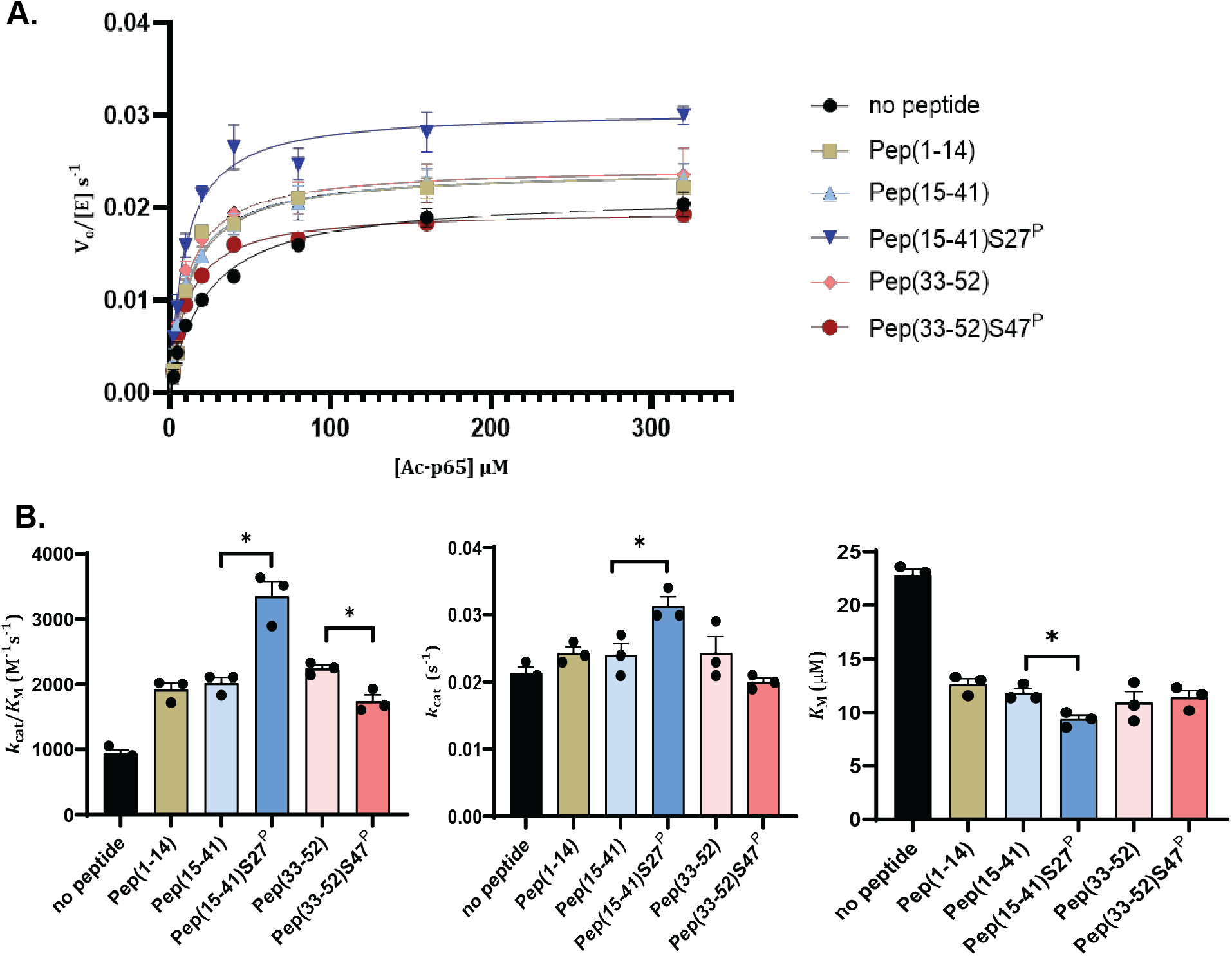
**A)** Enzyme kinetics curves for SIRT1-143 activity against Ac-p65 with and without the addition of various motif A-derived peptides. **B)** Michaelis-Menten parameters for SIRT1-143 activity with and without the addition of motif A-derived peptides. All kinetics data were collected in at least triplicates and fit with GraphPad Prism. The average and SEM are reported. ^*^ represent p-value < 0.05.

**Table 2.** Michaelis-Menten parameters for SIRT1-143 activity with and without motif A-derived peptides. All kinetics data were collected in at least triplicates and fit with GraphPad Prism. The average and SEM are reported.

| Additive | $k_{\text{cat}}$ (s<sup>-1</sup>) | $K_M$ (μM) | $k_{\text{cat}}/K_M$ (M<sup>-1</sup>s<sup>-1</sup>) |
| --- | --- | --- | --- |
| No peptide | 0.021 ± 0.001 | 23 ± 3 | 900 ± 100 |
| Pep (1-14) | 0.024 ± 0.001 | 13 ± 2 | 1900 ± 300 |
| Pep (15-41) | 0.025 ± 0.001 | 13 ± 2 | 1900 ± 300 |
| Pep (15-41) S27<sup>P</sup> | 0.030 ± 0.001 | 9 ± 1 | 3400 ± 500 |
| Pep (33-52) | 0.025 ± 0.001 | 11 ± 2 | 2200 ± 300 |
| Pep (33-52) S47<sup>P</sup> | 0.020 ± 0.0004 | 11 ± 1 | 1800 ± 200 |

### Molecular Dynamics Simulations Reveal the Effects of Phosphomimetics on Motif A Structure and Dynamics

Molecular dynamics simulations were conducted for WT motif A, motif A S27D, motif A S47D and motif A S27D S47D using the AlphaFold 3 generated structure predictions as a starting point. Averaging over the fully trajectories and across all residues, WT motif A and motif A S27D maintained similar helicity (26% and 27%, respectively), with motif A S47D averaging 33% helicity, and the double mutant motif A S27D S47D dropping substantially to 21% helical content. The location of the helices was relatively stable throughout the time steps for each construct, but were markedly different between the constructs (**Figure 5**). WT motif A and motif A S27D had stable helices within residues 3-9 (H1) and residues 15-25 (H2). While motif A S47D and motif A S27D S47D also exhibit a stable helix between residues 15-25, there is no persistent α-helix at the N-terminal, instead, both of these constructs form a helix between residues 31-34 (H3).

**Figure 5.**
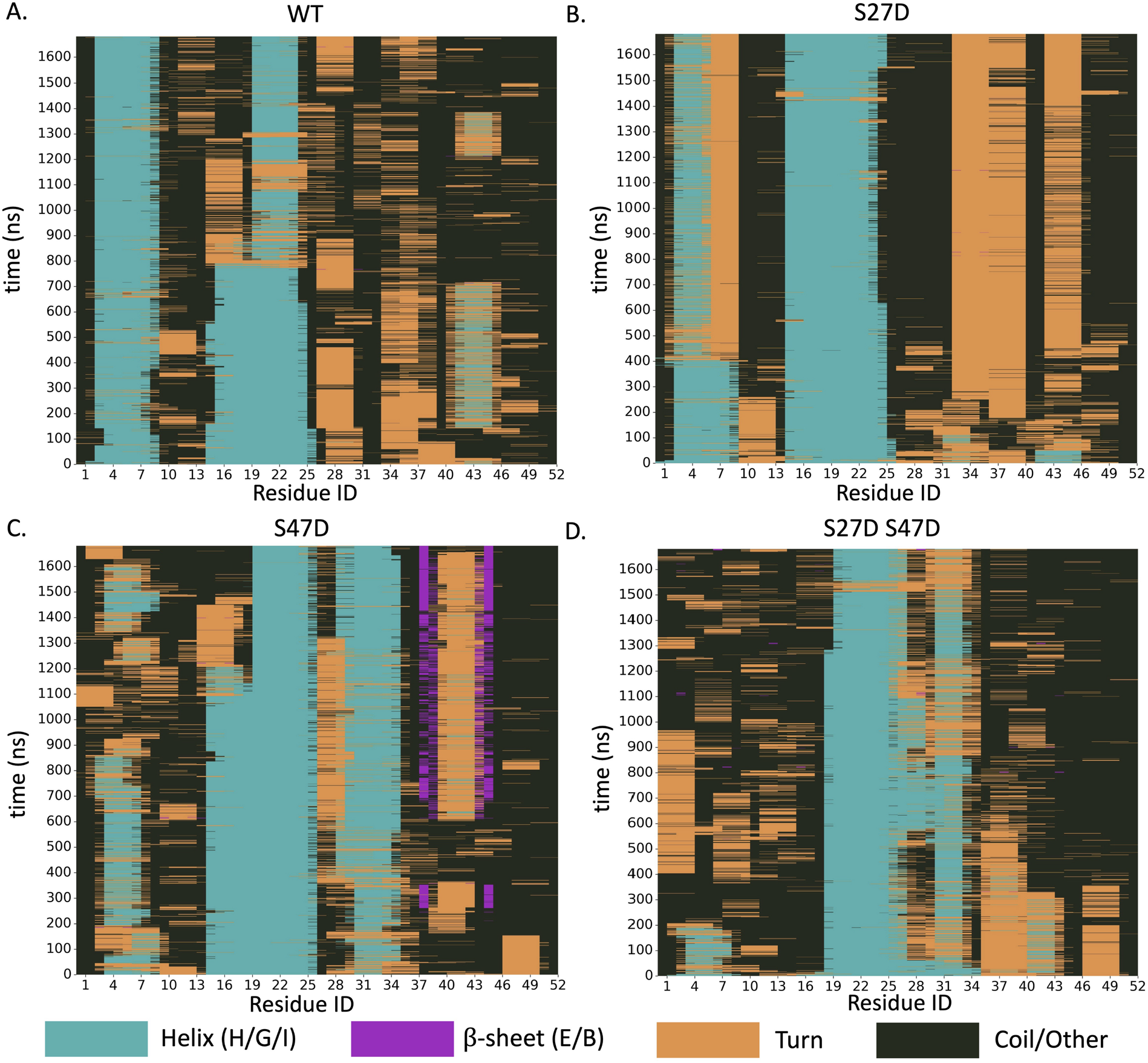
2D graph showing the secondary structure of each residue (x-axis) across all the simulated time steps (y-axis) for **A)** WT motif A, **B)** motif A S27D, **C)** motif A S47D and **D)** motif A S27D S47D.

This difference in structure was also reflected in the center of mass distance between H1 residues 3-9 and H3 residues 31-34. While the H1 and H3 residues are mainly 10 Å away from each other in the WT motif A and motif A S27D constructs, they were further apart (∼20 Å) in the motif A S47D construct. And the double mutant motif A S27D S47D construct exhibited populations at both distances (**Figure 6 A-B**). Furthermore, H2 and H3 seemed to be in a persistent “clamp” shape in motif A S47D and motif A S27D S47D. This shape might be stabilized by interactions between D21 and R34, which were extremely close in the motif A S47D construct and far away from each other in the motif A S27D construct (**Figure 6 C-D**).

**Figure 6.**
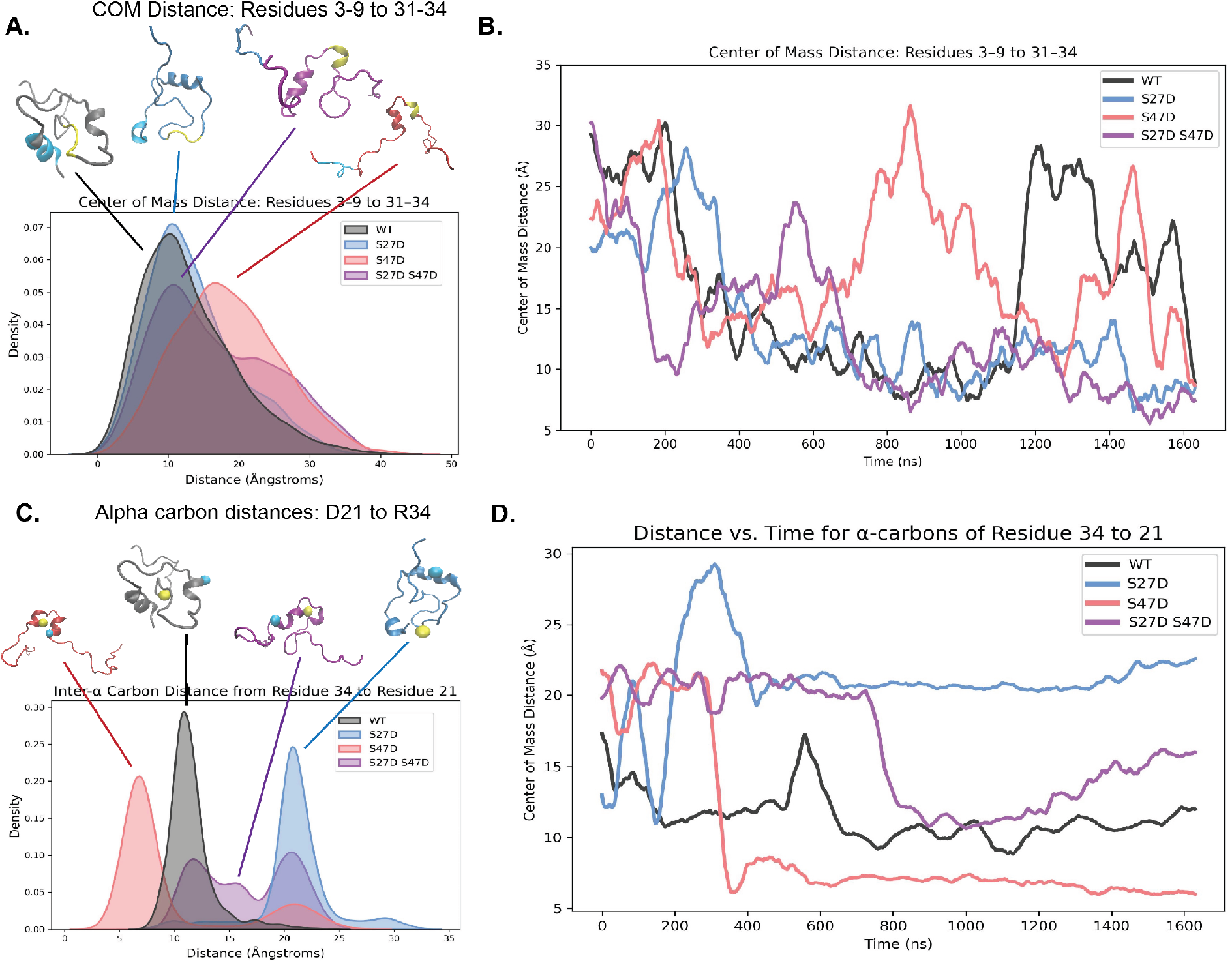
**A)** Distribution of center of mass distance between residues 3-9 and 31-34 for all four motif A constructs. Representative simulated structures are shown with residues 3-9 colored in cyan and 31-34 colored in yellow. **B)** Center of mass distance between residues 3-9 and 31-34 for all four motif A constructs over the simulation time frames (rolling mean smoothing applied with a 50 ps window for legibility). **C)** Distribution of inter-α carbon distance between D21 and R34 for all four motif A constructs. Representative simulated structures are shown with D21 colored in cyan and R34 colored in yellow. **D)** Inter-α carbon distance between D21 and R34 for all four motif A constructs over the simulation time frames (same smoothing as **B**.).

Further examination of the dynamics of the simulations revealed a remarkable difference between motif A S27D and the rest of the motif A constructs. Motif A S27D was significantly more rigid than the other constructs within the 1.7 μsec time scale of the simulation. This could be visualized as a small overall range of the radius of gyration as well as very low variance in the distance from the centroid by residue **(Figure 7A-B**). While not having significantly more secondary structure than the other motif A constructs, motif A S27D displayed much less movement, as demonstrated by the variance in motif A’s radius of gyration dropping precipitously from 4.1 Å^2^ in WT motif A to 0.08 Å^2^ in motif A S27D (Table SI 1). This is likely through the ionic interactions between a few key residues, namely D40 to R36, Q10 to R36, and D21 to R46, which seem to become very close to each other beyond the 500 ns time step (**Figure 7C**).

**Figure 7.**
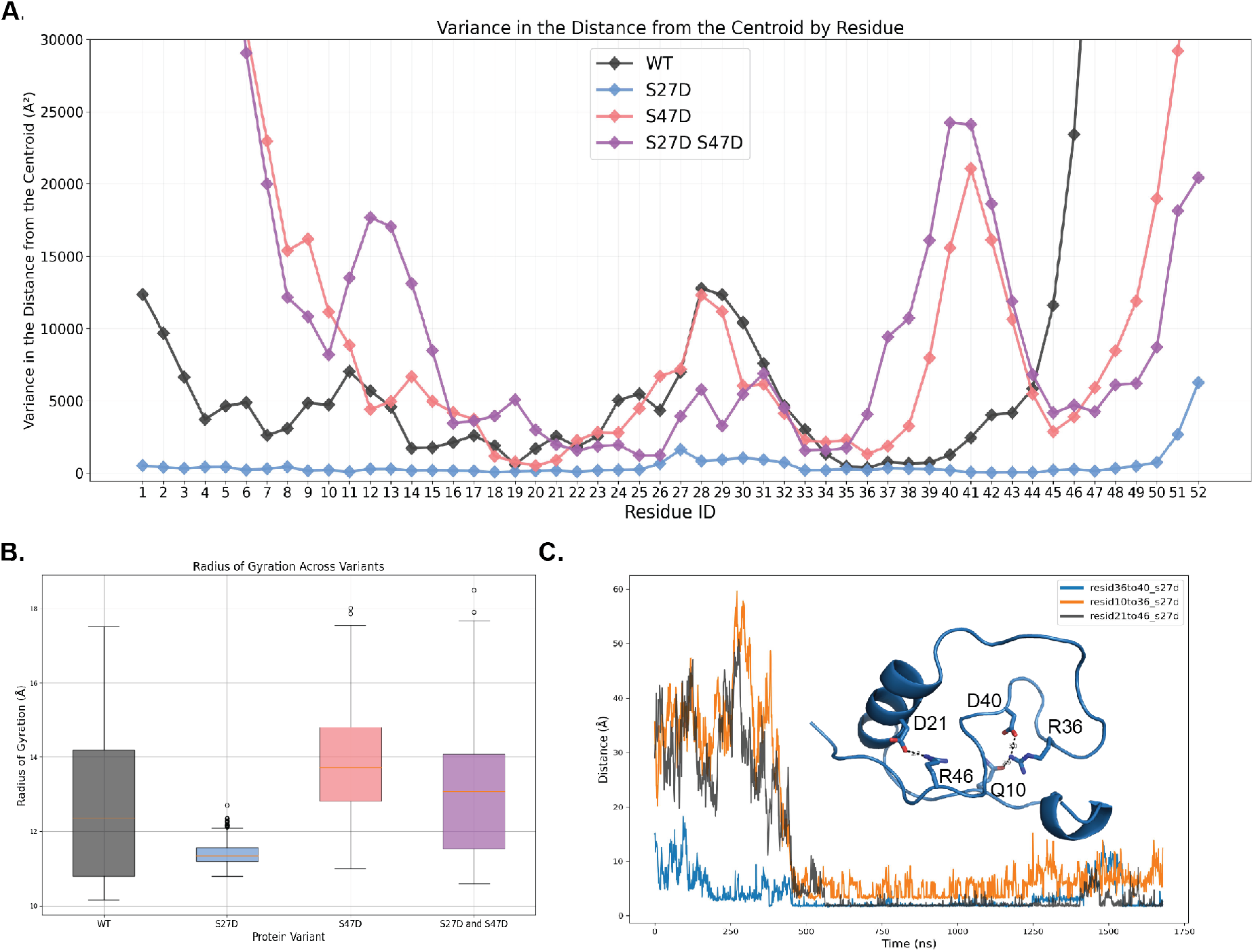
**A)** Variance in the distance from the centroid for each residue for all four motif A constructs. **B)** Radius of gyration across all simulation frames for all four motif A constructs. Thin red lines indicate the median, the colored boxes span the range from the 1st quartile to the 3rd quartile (i.e. the middle 50% of the values), whisker lines indicate the most extreme non-outlier values, and the hollow dots indicate outlier values. **C)** Distance between residues R36 and D40, Q10 and R36, and D21 and R46 over simulated time frame for motif A S27D. Insert shows representative simulated structure of motif A S27D with the residue in question represented in stick.

## Discussion

In this study, we were able to confirm that motif A increases SIRT1 activity *in vitro*, which fills in a knowledge gap regarding the role of motif A in SIRT1 regulation, as until now this activation effect has only been observed in cells^15^. More importantly, we demonstrated that the phosphorylation state of motif A plays an integral role in its ability to regulate SIRT1 activity. WT motif A does not affect SIRT1 activity. However, phosphomimetic mutation at S27 significantly increases the activation effect of motif A, which is also recapitulated in the assays with motif A-derived peptides containing authentic phosphorylation. The change in *K*_M_ is the main driving force for this activation effect, suggesting motif A S27D is able to alter the conformational dynamics of SIRT1 in a way that enhances its substrate recognition. This is consistent with physiological studies showing S27 phosphorylation is connected to protein activity and stability in the cell^21,22^. The effect of S47 phosphorylation on motif A’s regulation ability, however, is not entirely clear. Motif A S47D also activates SIRT1, albeit to a lesser extent than motif A S27D. However, this activation is achieved through an increase in *k*_cat_, the catalytic rate, suggesting that it has a different mode of activation than motif A S27D. This, however, is not recapitulated using the motif A-derived peptides, where phosphorylation at S47 doesn’t have a significant effect on the peptide’s ability to activate SIRT1. Previous studies have suggested that phosphorylation at S47 serves as a regulation for SIRT1 cellular localization^20^, hence it is reasonable to hypothesize that this phosphorylation event does not play a significant role in altering SIRT1 activity levels. Intriguingly, when both serines are mutated to aspartates, the motif A S27D S47D construct no longer has regulatory effects on SIRT1, similar to what we observe with WT motif A. This could possibly serve as a “down” regulator, where phosphorylating an extra serine can turn off the activation effects of motif A.

Molecular dynamics simulations of the motif A constructs provide possible clues to why motif A S27D interacts with SIRT1-143 differently from the rest of the constructs. First of all, the phosphomimetic mutation at S47 seems to shift motif A into a different secondary structure conformation, resulting in motif A S47D and motif A S27D S47D taking on different configurations from WT motif A and motif A S27D. This difference in structural landscape would very likely change the binding interface between motif A and the rest of SIRT1. Secondly, while WT motif A and motif A S27D have similar secondary structures, motif A S27D is shown to be significantly more rigid. This rigidity, likely aided by strong salt bridges between a few charged residues, might stabilize the interaction between motif A S27D and the rest of SIRT1 compared to WT motif A. This would explain why motif A S27D is also slightly more resistant to proteolysis, likely by being more rigid and less accessible to trypsin digestion. More importantly, this might be the reason why only motif A S27D has significant activating effects on SIRT1-143.

Taken together, our findings provide additional molecular insight into the regulation of SIRT1 within the cell, specifically focusing on how phosphorylation events within the intrinsically disordered motif A region could be used to fine-tune the activity levels of SIRT1. One key result of this study is the clear correlation between the rigidity of motif A S27D measured in the MD simulations and its experimentally observed activation of SIRT1-143 via its modulation of *K*_M_. This supports the hypothesis that phosphorylation of S27 serves as a molecular switch to activate SIRT1 by stabilizing the configuration of motif A that is able to bind to the rest of SIRT1.

## ASSOCIATED CONTENT

### Accession Codes

hSIRT1: Q96EB6

### Supporting Information

The following files are available free of charge. Additional Experimental Methods and Supplementary Figures (PDF)

## AUTHOR INFORMATION

### Author Contributions

The manuscript was written through contributions of all authors. All authors have given approval to the final version of the manuscript.

### Funding Sources

This work was supported by San José State University, NIH R16GM149377, and NIH R16GM150706. Instrumentation used for peptide synthesis was funded by NSF-1919817. Circular Dichroism instrumentation was funded by NIH S10GM154297.

## ACKNOWLEDGMENTS

We would like to thank Prof. Ruiming Xu, Prof. Jorge Escalante and Prof. Carol Fierke for providing plasmids for SIRT1-143, MBP-pncA and the SUMO protease.

## ABBREVIATIONS

CD: Circular Dichroism
MoRF: molecular recognition features

## Notes

### Competing Interest Statement

The authors have declared no competing interest.

