## Supplementary material for "The Effects of Phosphorylation on the Structure and Function of Motif A, an Intrinsically Disordered Region within SIRT1": SI

**Materials and Methods**

*Cloning of Motif A Phosphomimetic Constructs*

pET-28a-His_6_-TEV-SIRT1(1-52) plasmid was used for generating phosphomimetic mutations at S27, S47 and both S27 and S47. Site-directed mutagenesis (SDM) was performed using the New England Biolabs Q5 SDM kit (Ipswitch, MA). The primers used are shown in the table below. PCR products were transformed into NEB-10ꞵ cells and mutations were confirmed by sequencing.

Table S1. Primers used in site-directed mutagenesis

| S27D mutation | Forward Primer | 5’-GCGGCGAGCGATCCGGCGGG-3’ |
| --- | --- | --- |
|  | Reverse Primer | 5’-CTCACGGTCCGCACCCGCCG-3’ |
| S47D mutation | Forward Primer | 5’-TCTGGAGCGCGATCCGGTGAAC-3’ |
|  | Reverse Primer | 5’-CCCGGACCATCACGACGCGGACG-3’ |

*Expression and Purification of Motif A Constructs*

The pET-28a-His_6_-TEV-SIRT1(1-52) plasmids containing N-terminal His_6_ tagged SIRT1(1-52) constructs were transformed into BL21 (DE3) *E. coli* cells. A single colony was picked and grown in 20 mL Luria Broth containing 50µg/mL Kanamycin and incubated for 12 hours at 37°C, shaking at 220rpm. The overnight cultures were used to inoculate 1 L of TB in a 1:50 v/v ratio and were left shaking at 220rpm, 37°C until cells reached an optical density of 0.6-0.8 at 600 nm. Protein expression was induced with isopropyl β-d-1-thiogalactopyranoside (IPTG) to a final concentration of 0.1 mM. Cells were grown at 16°C for 20 hours and then centrifuged at 4500 rpm for 20 minutes, and dry pellets were stored at -80°C.

For purification, cell pellets were resuspended in lysis buffer (50mM TRIS pH 8.0, 150mM NaCl, 10% glycerol, 3mM 2-mercaptoethanol, 100µM PMSF), lysed by sonication, and clarified by centrifugation at 12000 rpm for 20 minutes. Soluble lysate was incubated with Ni-NTA resin and washed with lysis buffer. Protein was eluted in 50mM TRIS pH 8.0, 150mM NaCl, 10% glycerol, 3mM 2-mercaptoethanol in a gradient of 125 to 500mM imidazole. TEV protease was added to elution fractions to cleave His_6_ tag while protein was concurrently dialyzed into 2L storage buffer (50mM TRIS pH 8.0, 150mM NaCl, 10% glycerol, 1mM DTT) for 12 hours at 4°C. To further purify, the cleaved proteins were incubated with Ni-NTA and collected in flow through and wash fractions. Fractions were dialyzed into storage buffer at 4°C for 12 hours, and protein purity was verified by SDS-PAGE to be more than 85% pure. Samples were concentrated by centrifugation at 4000 g, the concentration of protein determined by Bradford assay, and flash frozen in liquid nitrogen, and stored at -80°C.

*Expression and Purification of SIRT1-143*

A truncated version of SIRT1 comprised of residues 143-500 and 641-665 termed SIRT1-143 was used for activity assays^1^. The plasmid of His_6_-SUMO-SIRT1-143, pET28-smt3-hSIRT1-143 was used to express SIRT1-143 in *E. coli* cells as described above. For purification, cell pellets were resuspended in a lysis buffer and lysed by sonication. Lysates were clarified by centrifugation at 12000 rpm for 20 minutes. Soluble lysate was incubated with Ni-NTA resin and washed in lysis buffer. Protein was eluted in 50mM TRIS pH 8.0, 150mM NaCl, 10% glycerol, 3mM 2-mercaptoethanol in a gradient of 125 to 500mM imidazole. Elution fractions were incubated with Ulp1 (SUMO protease) to cleave solubility tag while concurrently dialyzed into 2L storage buffer for 12 hours at 4°C. Subsequently, SIRT-143 was purified by size exclusion chromatography (Hiprep 16/20 Sephacryl S-160) on the ÄKTA™ Start FPLC system and eluted into storage buffer. Protein purity was verified by SDS-PAGE to be more than 85% pure. Samples were concentrated by centrifugation at 4000 g, the concentration of protein determined by Bradford assay, and flash frozen in liquid nitrogen, and stored at -80°C.

*Peptides Derived from motif A (SIRT1(1-52))*

Five different peptides spanning the first 52 residues of SIRT1, some with the phosphorylated S27 or S47 residues, were obtained. The names and sequences are listed in the following table. Pep(15-41) and Pep(15-41)S27^P^ were synthesized by the Hawk lab at Grand Valley State University. Pep(1-14), Pep(33-52) and Pep(33-52)S47^P^ were purchased from Elim Biopharm (Hayward, CA). All peptides were prepared to 95% purity as confirmed by HPLC.

Table S2. Peptide sequences

| *Peptide Name* | *Sequence* |
| --- | --- |
| Pep(1-14) | MADEAALALQPGGS |
| Pep(15-41) | PSAAGADREAASSPAGEPLRKRPRRDG |
| Pep(15-41)S27^P^ | PSAAGADREAASS^P^PAGEPLRKRPRRDG |
| Pep(33-52) | LRKRPRRDGPGLERSPGEPG |
| Pep(33-52)S47^P^ | LRKRPRRDGPGLERS^P^PGEPG |

*Peptide Synthesis*

Peptides were synthesized manually via Fmoc chemistry on a 0.025 mmol scale on NovaSyn TGR resin. For both coupling and deprotection, the resin and solution were bubbled gently with nitrogen to mix. Each coupling was carried out for one hour in DMF using four equivalents each of amino acid, 1-hydroxybenzotriazole (HOBt), and N,N,N′,N′-Tetramethyl-O-(1H-benzotriazol-1-yl)uronium hexafluorophosphate (HBTU), and 8 equivalents of DIEA. The Fmoc group was removed with a solution of 20% piperidine in DMF for 20 minutes. The resin was washed three times with DMF, three times with dichloromethane, and again three times with DMF between each step. Each reaction was checked for completeness with the ninhydrin test. Ser27 was phosphorylated through introduction of Fmoc-Ser-PO(OBzl)OH-OH. Attempts to synthesize phosphorylated SIRT115-41 through standard techniques were unsuccessful, so a kink was introduced into the peptide chain by incorporating Fmoc-Ala-(Dmb)Gly-OH dipeptide in place of Ala18 and Gly19. Incorporating the DMB-protected dipeptide resulted in the desired peptide after cleavage. After synthesis was complete, both peptides were cleaved from the resin using 95% TFA, 2.5% triisopropylsilane, and 2.5% water. Cleavages were carried out for 1.5 hours, after which the beads were filtered out of the solution. Excess TFA was evaporated under a stream of nitrogen and then cold ether was added to precipitate the peptide. The peptide was isolated through centrifugation and then dried under a stream of nitrogen gas. Peptides were purified via HPLC using a water-acetonitrile gradient where both solvents contained 0.1% TFA. Solvent was removed through lyophilization. Masses (Table S3) were confirmed using ESI mass spectrometry.

Table S3 Peptide mass spectrometry results

| Peptide | Expected Mass | m/z from ESI |
| --- | --- | --- |
| PSAAGADREAASSPAGEPLRKRPRRDG-NH_2_ | 2773.4 | M^+3^ 925.9  M^+4^ 694.7  M^+5^ 555.9 |
| PSAAGADREAASS_phos_PAGEPLRKRPRRDG-NH_2_ | 2853.4 | M^+3^ 952.7  M^+4^ 714.7  M^+5^ 572.2 |

**Supplemental Figures and Tables**

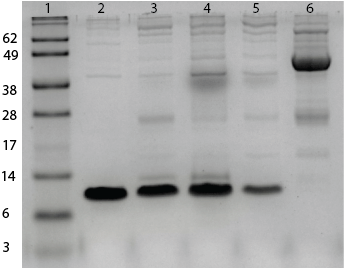

**Figure SI 1.** SDS-PAGE gel showing the purity of all purified proteins. Lanes are as follows from left to right: ladder, WT motif A (6.8 kDa), motif A S27D (6.8 kDa), motif A S47D (6.8 kDa), motif A S27D S47D (6.8 kDa), and SIRT1-143 (45 kDa).

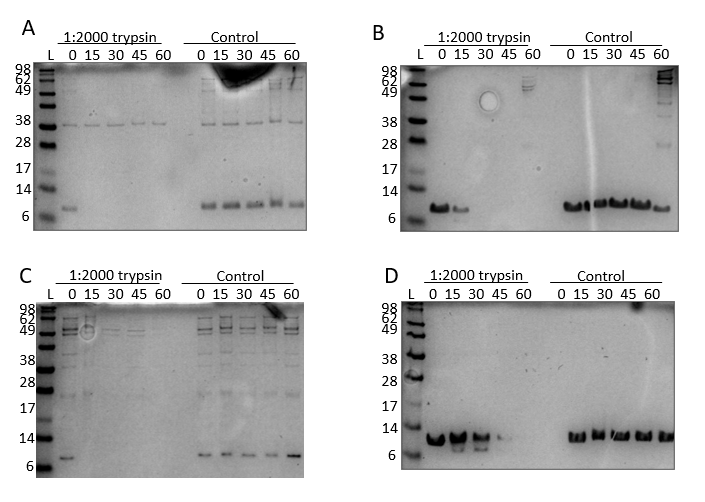

**Figure SI 2.** Commassie SDS-PAGE gels of limited proteolysis experiments of A) WT motif A, B) motif A S27D, C) motif A S47D and D) motif A S27D S47D. Control experiments were motif A constructs with no trypsin added, sampled and treated similarly over the same time frame.

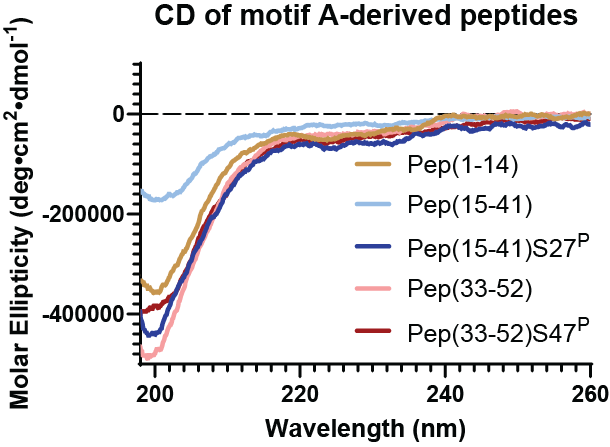

**Figure SI 3.** CD spectra of motif A-derived peptide fragments. All spectra exhibited random coil properties.

**Table SI 1.** Mean and variance of the radius of gyration of the 4 motif A variants over the course of their trajectories.

|  | **WT motif A** | **Motif A S27D** | **Motif A S47D** | **Motif A S27D S47D** |
| --- | --- | --- | --- | --- |
| Mean (Å) | 12.73 | 11.40 | 13.89 | 13.03 |
| Variance (Å^2^) | 4.11 | 0.08 | 1.85 | 2.36 |
